# Prioritizing Candidate eQTL Causal Genes in Arabidopsis using Random Forests

**DOI:** 10.1101/2022.04.06.487194

**Authors:** Margi Hartanto, Asif Ahmed Sami, Dick de Ridder, Harm Nijveen

## Abstract

Expression quantitative trait locus (eQTL) mapping has been widely used to study the genetic regulation of gene expression in *Arabidopsis thaliana*. As a result, a large amount of eQTL data has been generated for this model plant; however, only a few causal eQTL genes have been identified, and experimental validation is costly and laborious. A prioritization method could help speed up the identification of causal eQTL genes. This study extends the machine-learning-based QTG-Finder2 method for prioritizing candidate causal genes in phenotype QTLs to be used for eQTLs by adding gene structure, protein interaction, and gene expression. Independent validation shows that the new algorithm can prioritize sixteen out of twenty-five potential eQTL causal genes within the 20% rank percentile. Several new features are important in prioritizing causal eQTL genes, including the number of protein-protein interactions, unique domains, and introns. Overall, this study provides a foundation for developing computational methods to prioritize candidate eQTL causal genes. The prediction of all genes is available in the AraQTL workbench (https://www.bioinformatics.nl/AraQTL/) to support the identification of gene expression regulators in Arabidopsis.

## INTRODUCTION

One of the main objectives of genetic research is to link traits to genotypic variation. However, the path from genetics to observable traits is not straightforward; instead, it goes through a network of interconnecting intermediate phenotypes, such as gene expression, protein levels, and metabolite levels (Civelek and Lusis 2013). Studying the effect of the genetic perturbation on these intermediate phenotypes could improve our understanding of how a trait is regulated. Following recent advances in omics technology, the effect of multiple genetic perturbations can now be studied in a single experiment using linkage mapping or association studies. One example is genetical genomics, where variation in transcript levels is statistically associated with genetic variation in a population (Jansen and Nap 2001) to find so-called expression quantitative trait loci (eQTLs).

A mapped eQTL can be categorized as *cis* or *trans* based on its location relative to the affected gene. *Cis*-eQTLs are mapped close to the gene and are assumed to arise due to sequence polymorphisms in or near the gene itself, for instance, in *cis*-regulatory elements (*e*.*g*., the promoter). In contrast, *trans*-eQTLs are mapped far away from the target gene and emerge due to polymorphisms in *trans*-acting factors (*e*.*g*., transcription factors) called expression quantitative trait genes or eQTGs (Rockman and Kruglyak 2006; Brem et al. 2002). However, a *trans*-eQTL typically spans a large genomic region with hundreds of candidate eQTGs. Experimental fine mapping to narrow down the region (*e*.*g*., in Eshed and Zamir 1995) is costly and laborious. As a result, only a few causal genes have been identified in the thousands of eQTLs that have been mapped for *Arabidopsis thaliana*, using different populations and experimental conditions (Keurentjes et al. 2007; West et al. 2007; Cubillos et al. 2012; Snoek et al. 2012; Lowry et al. 2013; Hartanto et al. 2020). As an *in silico* alternative, a prioritization method can help to limit the number of candidate eQTGs for further validation.

Several network-based methods have been used to find eQTGs (*e*.*g*., in Keurentjes et al. 2007; Jimenez-Gomez et al. 2010; Hartanto et al. 2020). These methods primarily aim to find master regulator(s) at loci where *trans*-eQTLs for many genes are collocated, known as eQTL hotspots (Breitling et al. 2008). In general, these methods utilize a coexpression network built using genes having an eQTL on the hotspot (called *targets*) and genes located in the hotspot (called *candidate eQTGs*). Candidates are then usually prioritized based on a network centrality measure, such as degree or closeness centrality (Serin et al. 2016). Several candidate eQTGs have been identified in this way, for example, *GIGANTEA* (Keurentjes et al. 2007), *ELF3* (Jimenez-Gomez et al. 2010), *ICE1*, and *DEWAX* (Hartanto et al. 2020). This approach, unfortunately, only works for eQTL hotspots, not for regions that only have a small number of eQTLs. Another limitation is the sole reliance on coexpression data: given the complexity of gene expression regulation, the expression of the regulator is not necessarily correlated to that of its targets, particularly in eukaryotes (Marbach et al. 2012; Lelli et al. 2012). Therefore, additional data sources should be considered to capture possible interactions between the regulator and its target.

Previously, a machine-learning-based method, QTG-Finder, was developed to prioritize candidate genes for phenotype QTLs in Arabidopsis (Lin et al. 2019). This method used features derived from various gene properties, such as paralog copy number, gene ontology, and the number of SNPs, to rank the candidate genes in the QTL interval. The model could recall 64% of Arabidopsis QTGs when the top 20% ranked genes were considered. Further development of this method led to QTG-Finder2, which used orthology information and allowed for gene prioritization in species with no or few known QTGs (Lin et al. 2020). We were curious about the capability of this algorithm to prioritize eQTGs, given that some QTGs are involved in gene expression regulation, for example, *ELF3* (Jimenez-Gomez et al. 2010), *ERECTA* (Terpstra et al. 2010), *FRI* (Lowry et al. 2013), *MAM1* (Jansen et al. 2009), and *AOP2* (Jansen et al. 2009).

We propose eQTG-Finder, an extended version of QTG-Finder2 for eQTG prioritization, and apply the new algorithm to prioritize eQTGs in Arabidopsis. eQTG-Finder contains twelve new features based on protein-protein interaction, gene structure, and expression variation. These features significantly improve model performance, which is underscored by a feature importance analysis. We demonstrate the efficacy of this algorithm in prioritizing eQTGs using an independent test set. Finally, we use the new model to predict all Arabidopsis genes and make these available in our Arabidopsis eQTL analysis platform AraQTL (https://www.bioinformatics.nl/AraQTL/)(Nijveen et al. 2017) to help identify gene expression regulators.

## MATERIALS AND METHODS

QTG-Finder2 was developed for prioritizing causal phenotype QTL genes (QTG) in Arabidopsis (Lin et al. 2020). This algorithm consists of 5,000 Random Forest classifiers (Ho 1998) trained using known QTGs and Arabidopsis orthologs of QTGs from other species as positives and other genes as negatives. QTG-Finder2 prioritizes candidate genes based on features generated from polymorphism data, functional annotation, co-function networks, and paralog copy numbers. Our method extends QTG-Finder2 with new features, and we train the resulting model using the same sets of positive and negative genes. We evaluate the performance in prioritizing candidate causal eQTL genes (eQTGs) in Arabidopsis.

### New features

We generate and include twelve new features in addition to the ones already used by QTG-Finder2. These new features are based on protein-protein interactions, gene expression, and gene/protein structure.

1. Protein-protein interaction feature Genes can be associated with other genes, for instance, because the encoded proteins participate in the same pathway or are mentioned in the same publication. The number of such interactions a gene has could measure its propensity to be an eQTL causal gene. We generate a network-based feature using Arabidopsis protein-protein interaction (PPI) data from STRING-DB (Szklarczyk et al. 2019). The data were downloaded from the download page of STRING-DB version 11 (https://string-db.org/cgi/download). We only keep high-confident interactions by removing those with STRING scores below 700. We count the number of interactions of each Arabidopsis gene as a feature.
2. Gene expression features The consequence of genetic variation in causal genes might be detected as early as in gene expression variability. We, therefore, generate features based on gene expression variation. We use the standard deviation of expression levels across different tissues from CoNekT (http://www.evorepro.plant.tools/) (Julca et al. 2020). We also use the average and standard deviation of *Arabidopsis thaliana* Columbia ecotype expression data from different samples as features. These data were retrieved from the Athrna-database (http://ipf.sustc.edu.cn/pub/athrna/) (Zhang et al. 2020).
3. Structural features The structure of causal genes and encoded proteins might differ from the other genes. Therefore, we generate structural features: the numbers of introns, splice variants, total protein domains, unique protein domains, and splice variants per gene. Data were retrieved from https://www.arabidopsis.org/ (accessed May 2021). The number of introns and splice variants are counted in TAIR10’s BLAST datasets. The other two features are generated from all.domains.txt by counting each Arabidopsis gene’s total number of domains and the number of unique domains.

### Hyperparameter tuning

Model evaluation is based on QTG-Finder (Lin et al. 2019) and QTG-Finder2 (Lin et al. 2020). Similar to QTG-Finder2, we use known QTGs and Arabidopsis orthologs of QTGs found in other species as positives and other genes as negatives. We use hyperparameter tuning to determine the best parameter combination (the number of trees, minimal samples split, and maximum number of features) using grid search and assess the area under the curve (AUC) of the receiver characteristics operator (ROC) curve in an extended version of the 5-fold cross-validation framework. In this framework, the positives are randomly re-split into a training and validation set in a 4:1 ratio iteratively. Next, each set is combined with randomly selected negatives. The ratio of positives and negatives is an optimized hyperparameter. This splitting of positives is done 50 times, and for each positive set random selection of the negatives was conducted 50 times. This extensive procedure (2,500 evaluations) makes that positive co-occurs with all negative at least once with high probability. All machine-learning model training and testing in this study is performed using Python’s scikit-learn library version 1.0.2.

### Selection of candidate eQTL genes and independent validation of model performance

A list of candidate eQTGs in Arabidopsis is manually selected from the literature. These genes are categorized as confirmed/strong-candidate, hypothetical, or hypothetical-ortholog. Genes that have been through experimental validation or have strong evidence as eQTG are categorized into the confirmed/strong-candidate group, for example, *GIGANTEA* (Keurentjes et al. 2007; Snoek et al. 2012). Some confirmed/strong-candidate eQTGs are used as positive in QTG-Finder2, and we remove these from the positive instances to be used as validation genes. Meanwhile, genes that were not experimentally validated but are predicted to play a role as eQTG through *in silico* analysis (*e*.*g*., network analysis) are categorized as hypothetical, for example, *ICE1* and *DEWAX* (Hartanto et al. 2020). If a gene’s ortholog is considered an eQTG in another species, it is categorized as hypothetical-ortholog; for example, *NF-YC4* is found as an eQTG in potatoes (van Muijen et al. 2016). In total, this yields twenty-five candidate eQTGs in Arabidopsis: six confirmed/strong-candidate, four hypothetical, and fifteen hypothetical-ortholog genes (Supplementary Table 1). We ensure that these candidates are not used for hyperparameter tuning or cross-validation.

Independent validation is performed using the best combination of parameters (Supplementary Table 1). We train 5,000 Random Forest classifiers using all positives but different sets of negatives, with a positive: negative ratio of 1:200 to approximate the ratio of causal and non-causal genes in real eQTLs. The models are then applied to each candidate eQTG and other genes located 1 Mbp around it. These genes are ranked based on the average probability of being causal genes over 5,000 models.

### Feature importance analysis

Feature importance is determined using a leave-one-out analysis. Iteratively, each feature is removed from the dataset, and a model is trained using the reduced dataset. The AUC difference in the full model (with all features) and the reduced model is then calculated and used to indicate the feature importance. We use the previous cross-validation framework and the best parameters to measure the model performance in this analysis.

### Data analyses and code availability

Pairwise Pearson correlation coefficients between features are calculated using the Pandas (version 1.3.5) DataFrame.corr method in Python. Pearson Wilcoxon Rank Sum Test tests differences in the median between positive and negative genes for the twelve new features. The test is conducted in R using the base ‘wilcox.test’ function. Gene ontology enrichment analysis for the top and bottom 5% predicted causal genes is performed using TopGO in R (Alexa et al. 2006) using the algorithm’s default ‘weight01’ parameter, which is the mixture of ‘elim’ and ‘weight’ methods. The Python version used for the analyses is 3.8.12, and the R version is 4.0.2. The source code and data are available at https://git.wur.nl/harta003/eqtg-finder.

## RESULTS

The QTG-Finder2 algorithm could rank phenotype QTL causal genes higher than other genes in a cross-validation setting (AUC = 0.81) and recall 80% independent curated causal genes when the top 20% of genes in the QTL are considered (Lin et al. 2020). In this study, we extend QTG-Finder2 with a set of new features and evaluate its performance in prioritizing expression QTGs.

### New features improve causal gene prediction performance

To improve model performance and better tailor it fit for eQTG prioritization, we added twelve new features based on gene expression, structure, and protein-protein interactions in the QTG-Finder2 algorithm. Most new features only show a low to moderate correlation with the existing ones (Supplementary Figure 1), indicating that we add new information to the model. Figure 1 shows feature distributions for the causal genes as the positive class (55 known QTGs and 145 Arabidopsis ortholog of QTGs from other species) and the other genes in the genome as the negative class (n=26,970). For most features, the causal genes’ median value is significantly different from that of the other genes in the genome (see Supplementary Table 3). The expression of causal genes is more variable than that of other genes. Moreover, causal genes tend to have more and varied protein domains. Causal genes also have slightly more introns than other genes. These differences between the causal genes and the other genes in the genome provide a first indication of potential discriminating features for the machine learning model. We assess the performance of the model with and without new features using a cross-validation framework.

**Figure 1.**
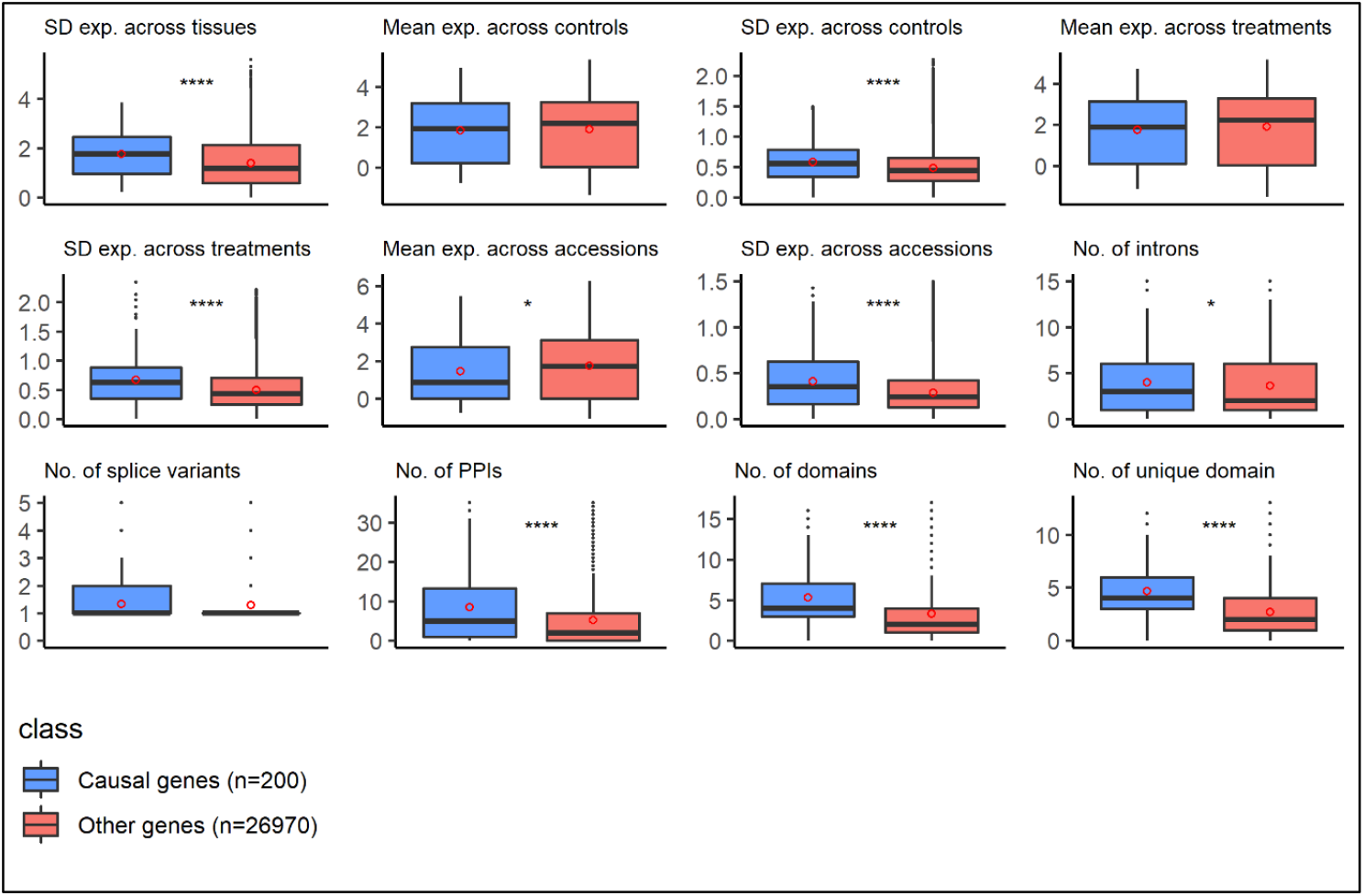
Distribution of twelve new features for known causal genes as the positive class (blue: n=200; 55 known QTGs and 145 orthologs of QTGs from other species) and the remaining genes in the genome as the negative class (red: n=26,970). Significance of differences in medians was assessed using the Wilcoxon Rank Sum Test (*: p <= 0.05; ****: p <= 0.0001). Red dots indicate means. SD = standard deviation. Exp. = gene expression. PPIs = protein-protein interactions.

To assess the contribution of new features to the model performance, we compare the area under the receiver-operating characteristic curve (AUC) between the original QTG-Finder2 with the extended model that we labeled eQTG-Finder, and for the extended model with the class labels permutated, as a control (Figure 2 left). The AUC was measured in an extended cross-validation setting over 2,500 different combinations of positive and negative gene sets. The results show that eQTG-Finder (AUC = 0.859 ± 0.008) performs better than QTG-Finder2 (AUC = 0.801 ± 0.01) and the control model (AUC = 0.502 ± 0.014). Adding new features thus allows the model to rank causal genes higher than the other genes. The next section analyzed model performance in prioritizing eQTG using selected candidate eQTGs.

**Figure 2.**
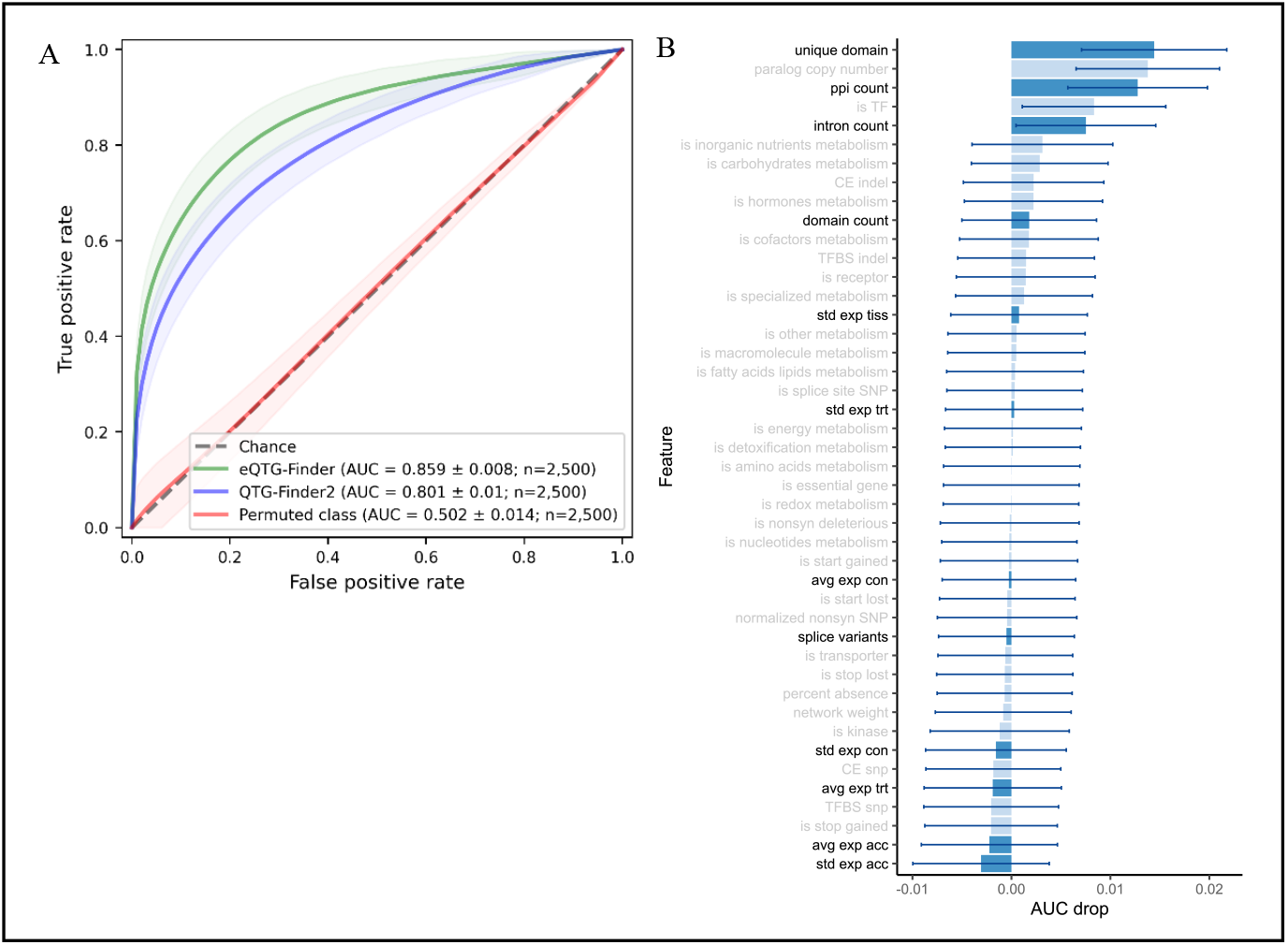
(A) Receiver operating characteristic (ROC) curves of the original QTG-Finder2 model (blue) and extended eQTG-Finder model (green), and eQTG-Finder trained with randomized class labels (red) as a control. Transparent areas indicate standard deviations over 2,500 repetitions. (B) Feature importance is measured using leave-one-out analysis. A positive AUC drop indicates that the removal of the feature reduces the model’s predictive capability. Feature names in bold and with dark blue bars indicate new features. Error bars indicate standard deviations over 2,500 repetitions.

To determine how the new features contribute to causal gene prediction, we calculate feature importance using a leave-one-out approach. Each feature is iteratively removed from the dataset, and the reduced model’s performance is compared to that of the model containing all features. The drop in AUC indicates a feature’s importance. A positive AUC drop means removing that feature decreases the model’s predictive capability. The result shows that four of the most important features in the model are the new ones: the number of unique domains, the PPI count, the intron count, and the domain count. However, the large standard deviation for the domain count AUC drop indicates that the contribution of this feature is not consistent over different samples of positive and negative sets.

### eQTG-Finder ranks most strong eQTG candidates better than QTG-Finder2

To evaluate eQTG prioritization performance, we again train the original QTG-Finder2 and the extended eQTG-Finder model and use them to rank selected potential eQTGs (Supplementary Table 1). Models are trained using all positives (known QTGs and Arabidopsis ortholog QTGs from other species). We repeated the training 5,000 times with different negative samples to select each negative gene at least once in training with high probability. These models rank each of the twenty-five potential eQTGs with their surrounding genes within a 2 Mbp window as a hypothetical eQTL region. These potential eQTGs are selected manually from the literature and grouped based on the evidence of being causal eQTL genes (see Methods for detail). Gene ranking is based on the average probability of a gene being causal, as predicted by the 5,000 models. We use the rank percentile to indicate the percentage of genes on the eQTL with higher ranks than the gene of interest (*i*.*e*., a rank percentile of 0.1 indicates that 10% of genes in the eQTL region rank higher than the gene of interest). We predefine cutoffs of 5%, 10%, and 20%, in each of which we compare recall between QTG-Finder2 and eQTG-Finder. These recalls for different cutoffs can be used by researchers to decide the proportion of top prioritized genes for further experimental validation.

The QTG-Finder2 model recalls 16%, 28%, and 52% of eQTG candidates if the top 5%, 10%, and 20% ranked genes are considered (Figure 3). With added features, eQTG-Finder ranks eQTGs slightly better with percentages of 36%, 52%, and 64% respectively. The eQTGs vary in their evidence of being causal genes (see Methods). Four out of six strong eQTG candidates (*AOP2, ERECTA, GIGANTEA*, and *MAM1*) rank within the 5% rank percentile by eQTG-Finder compared to only one (*ERECTA*) by QTG-Finder2. The other two strong candidates, *FRI* and *ELF3*, were ranked at the 10.2% and 61.2% percentile by eQTG-Finder. The ranks of sixteen genes are improved by eQTG-Finder, eight are worse, and one stays the same (Supplementary Table 4). The rank of four out of six strong eQTG candidates improves, with *GIGANTEA* one of the most drastic improvements, moving from 53,7.7% to 4.2%. On the other hand, the rank of *ERECTA* drops (0.4% to 2.8%) but remained falls in the 5% rank percentile. Both models rank another strong eQTG candidate *ELF3* poorly (at 44% rank percentile by QTG-Finder2 and 61.2% by eQTG-Finder). Despite the decent overall performance in eQTG prioritization, we notice that eQTG-Finder performance in prioritizing phenotype QTGs is still inconsistent. Using the initial independent validation set, only seven out of eleven QTGs are ranked within the 20% rank percentile by eQTG-Finder, compared to nine by QTG-Finder2 (Supplementary Figure 2).

**Figure 3.**
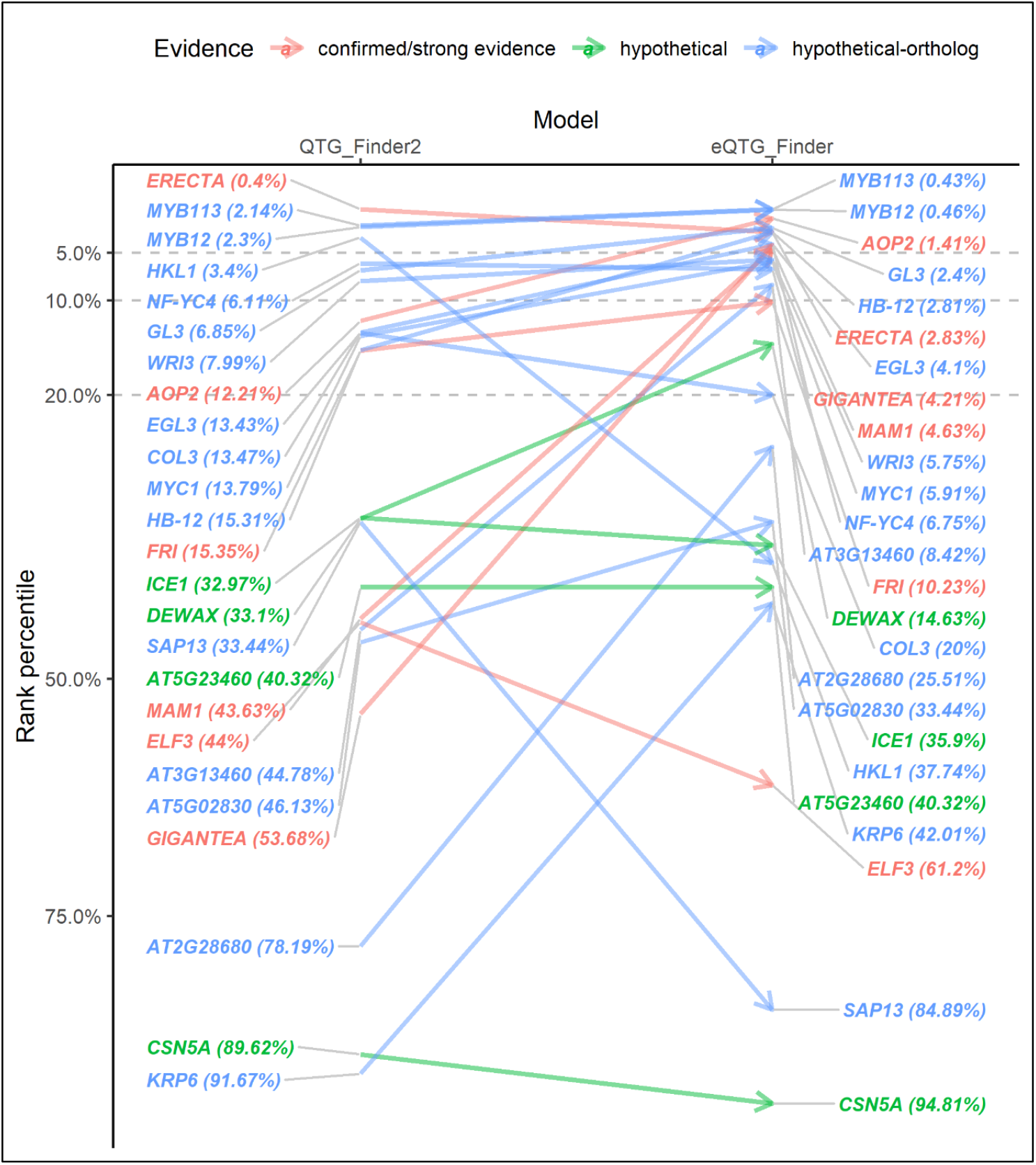
Rank percentile comparison of sixteen candidate eQTGs using the model with new features (eQTG-Finder) and the original model (QTG-Finder2).

To get an overview of eQTG-Finder predictions, we inspect the distribution of the average predicted probability of being causal for all Arabidopsis genes (Figure 4). This skewed towards a low value, with a median value of 0.007 (note that the *x*-axis of Figure 4 is on a log_10_ scale). Twenty-one of the twenty-five genes in the validation set have a predicted probability higher than the median. *ELF3* (probability=0.0045) is the only strong eQTG candidate with a predicted probability lower than the median. A Gene Ontology (GO) enrichment analysis shows that the top 5% genes in the distribution are significantly enriched (FDR p-value < 0.05) for 67 GO terms (Supplementary Figure 5), most of which are related to response to abiotic and biotic stresses, such as “defense response to bacterium”, “defense response to fungus”, and “response to wounding”. The term “regulation of transcription” is also enriched, suggesting that transcription factors are likely to be causal, consistent with the feature importance analysis result where is_TF is among the most important features. Meanwhile, the bottom 5% are not enriched for any term.

**Figure 4.**
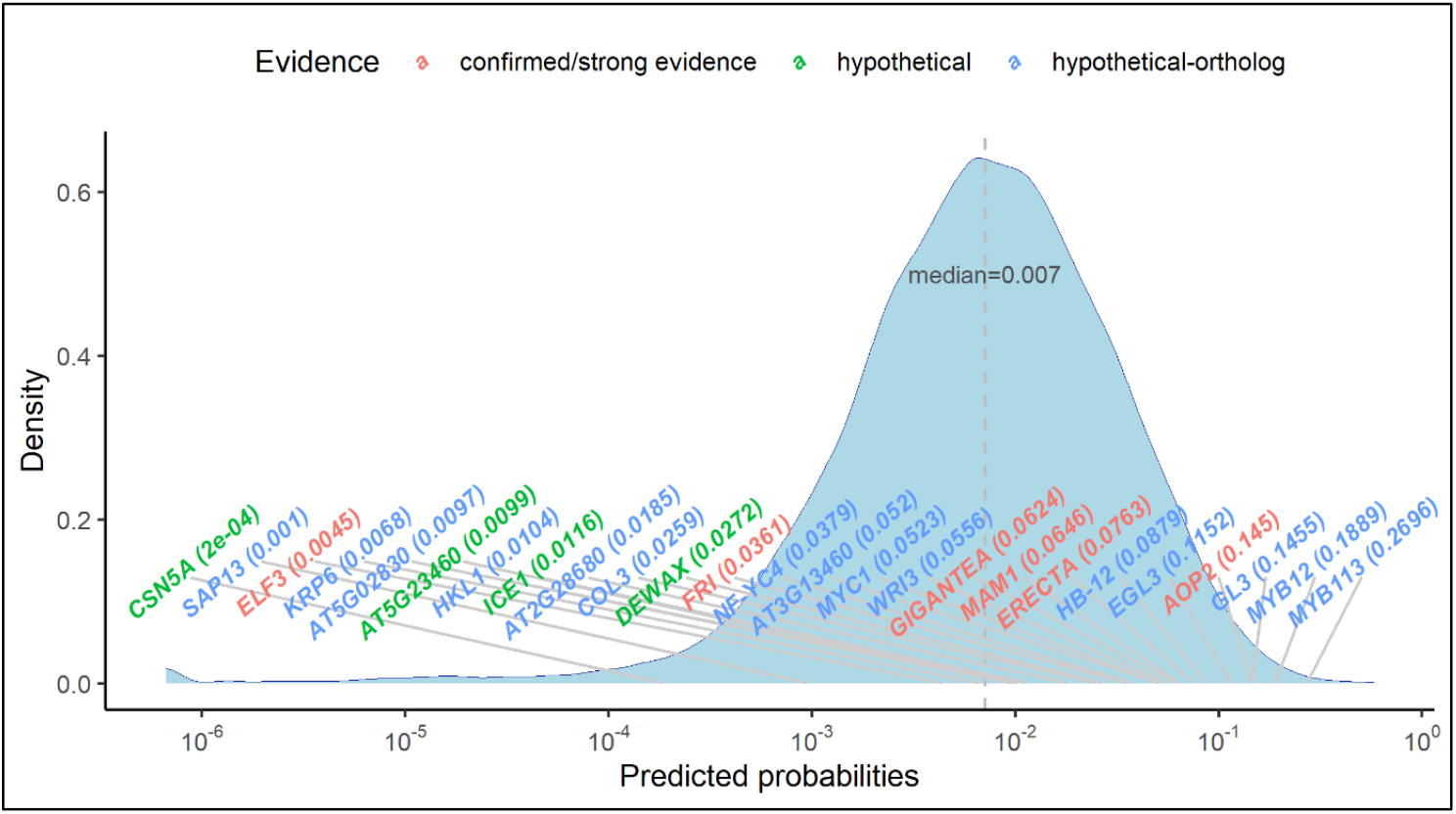
The density plot of probabilities of being causal predicted by eQTG-Finder for all Arabidopsis genes. Text labels point to the probability of the gene in the plot. The *x*-axis is on a log_10_ scale.

**FIGURE 5.**
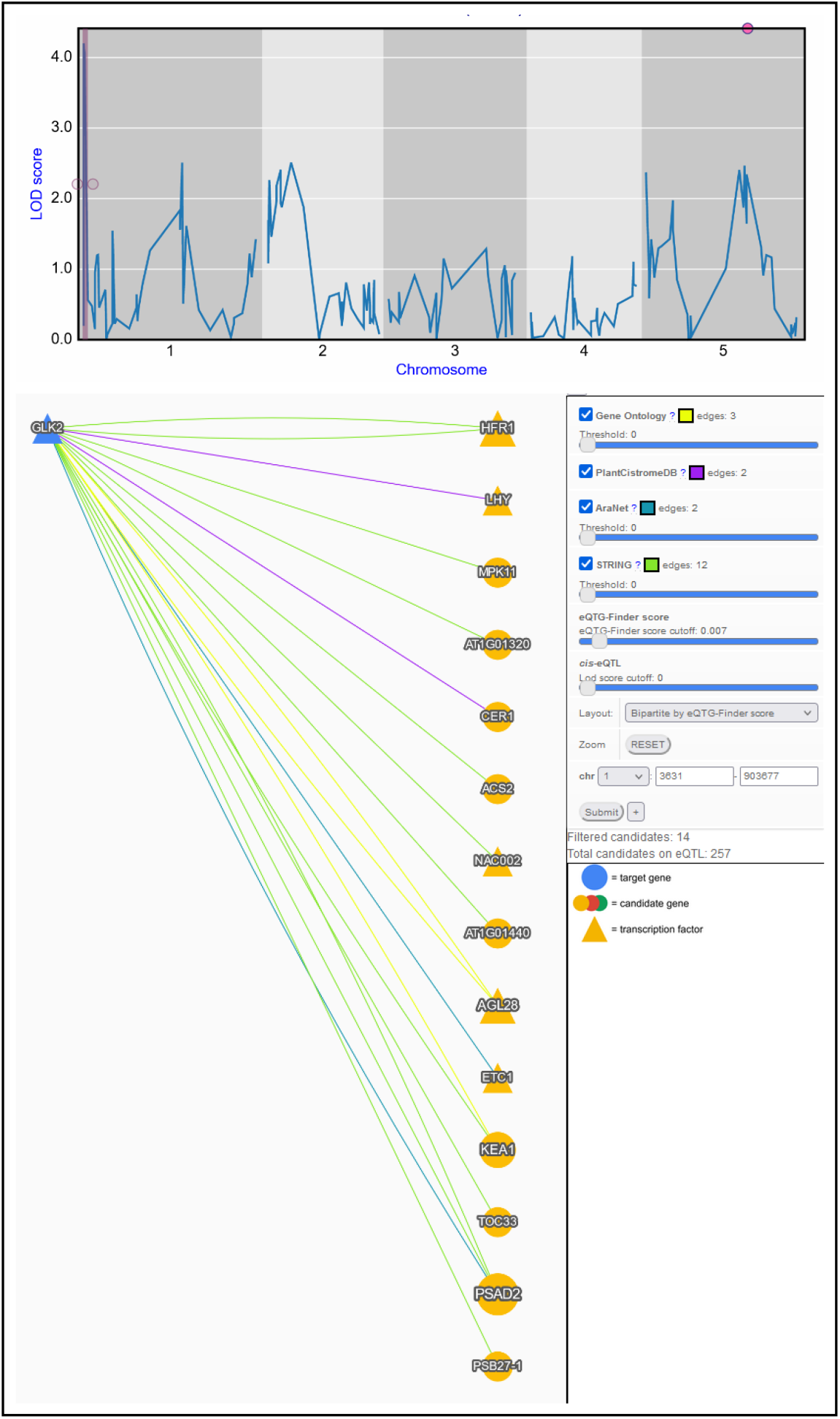
Prioritization of *GLK2* regulator using the eQTG-Finder score in AraQTL. (top) eQTL profile of *GLK2* from the Lowry et al. (2013) experiment. The eQTL region on chromosome 1 (shaded in pink) pinpoints the location of potential *GLK2* regulator(s). (bottom) Prior-knowledge network connecting *GLK2* (blue node) with candidate regulators (yellow nodes) based on prior knowledge data. Here, the eQTG-Finder score is used to order candidates based on their probability of being causal.

### eQTG-Finder is available in AraQTL to support new hypotheses on the gene expression regulation

To make eQTG-Finder results easily accessible for researchers, we include predicted probabilities of causality (herewith referred to as eQTG-Finder score) for all Arabidopsis genes in AraQTL, our Arabidopsis eQTL data workbench (Nijveen et al. 2017). Prioritizing genes using QTG-Finder2 is not straightforward as it requires users to prepare a list of candidate genes and command-line usage skills. Integrating the eQTG-Finder score in AraQTL facilitates users to interactively identify gene expression regulators. For example, we here discuss a case on predicting a new potential regulator for *GLK2* using the eQTG-Finder score and other interaction evidence in AraQTL. *GLK2* is a GARP nuclear transcription factor involved in light-controlled signaling (Waters et al. 2009). Liu et al. (2021) recently found that *HY5* is the regulator of *GLK2* based on the fact that *HY5* is a well-known regulatory switch for light signaling in literature. The same conclusion can also be derived using the Serin et al. (manuscript in preparation) eQTL experiment and prior knowledge data in AraQTL. Another approach to finding potential regulators of *GLK2* can be made in AraQTL using the eQTG-Finder score. In a Kas x Tsu eQTL experiment on leaf tissue (Lowry et al. 2013), *GLK2* has an eQTL on the beginning of chromosome 1, indicating the location of the potential regulator(s) (Figure 5, top). As many as 257 candidate regulatory genes are present in the eQTL (Figure 5, bottom). We can filter out weak candidates by constructing a network of *GLK2* connected to its potential regulators on the eQTL based on prior knowledge, such as protein-protein interaction and gene annotation (Hartanto *et al*., manuscript in preparation). Here, we threshold the eQTG-Finder score to remove weak candidates. Moreover, eQTG-Finder can prioritize the remaining fourteen genes by selecting the “Bipartite by eQTG-Finder score” network layout and ordering genes by their score. The result suggests some promising *GLK2* regulator candidates ranked at the top, for example, a transcription factor *LHY* in second place. Until now, *LHY* has not been reported to regulate *GLK2*. However, this gene is a promising *GLK2* regulator candidate as the network shows that it has a transcription factor binding site(s) on the *GLK2* promoter (O’Malley et al. 2016). Moreover, *LHY* is involved in light signaling (Joo et al. 2017; Kim et al. 2003). This example suggests that integrating the eQTG-Finder score in AraQTL can help infer new regulatory interactions.

## DISCUSSION

The concept of genetical genomics was first coined two decades ago (Jansen and Nap 2001), and numerous Arabidopsis eQTL data sets have been published since then (Nijveen et al. 2017). The aim of genetical genomics is to pinpoint genomic regions associated with gene expression variation (eQTL) and ultimately unravel genes involved in expression regulation. However, identifying causal genes (eQTGs) is difficult because of the often large genomic regions they span, regularly harboring dozens or even hundreds of candidates. The regions can be narrowed down by experimental fine-mapping (Eshed and Zamir 1995), and the remaining candidate genes can then be validated using functional genomics methods (*e*.*g*., using CRISPR-Cas9-mediated deletions as in Evans and Andersen 2020). However, performing these experiments for thousands of eQTLs is very costly. Using genomics and annotation data, a computational prioritization method can help identify candidate eQTGs. This study extends an existing machine-learning algorithm, QTG-Finder2, to address this issue and evaluates its performance for prioritizing eQTG. eQTG-Finder outperforms its predecessor in a cross-validation setting and independent validation test. We make eQTG-Finder scores available in AraQTL to help researchers interactively identify key regulators.

The key improvement of eQTG-Finder lies in the inclusion of twelve new features based on gene expression, structure, and interactions. Given the complexity of the resulting model, it is not straightforward to assess how these features improve eQTG-Finder in gene prioritization (Petch et al. 2022). We calculated the contribution of each feature in the model using a leave-one-out feature importance analysis (see Materials and Methods). This showed that the number of unique protein domains, the number of protein-protein interactions (PPI), and the number of introns are in the top five most contributing features in the model. We showed that known causal genes tend to have more domains, protein-protein interaction partners, and introns than other genes (Figure 1). These new features may provide insight into what distinguishes causal and non-causal genes. For instance, since protein domains determine protein functions (Vogel et al. 2004; Enright and Ouzounis 2001), the presence of multiple domains in a causal gene could indicate involvement in a wide range of biological functions. The diverse functions of causal genes could also be reflected in their larger number of protein-protein interaction partners than non-causal as genes perform their function in concert with other genes (Ito et al. 2001). The number of introns reflects the number of exons in a gene. Several studies demonstrated that exons play a role in the evolution of domain architectures through exon-shuffling, leading to new combinations of domains with new functions.

Variation in phenotype can be traced back to variation in gene expression (Skelly et al. 2009; Albert and Kruglyak 2015). For this reason, we included features based on the standard deviation (SD) of gene expression across different Arabidopsis accessions and conditions. Even though the medians between causal and other genes are significantly different (Figure 1), features based on SD of expression have low importance in the model. A study showed that correlations between features decrease the importance to zero (Gregorutti et al. 2016). Given that three SD features are highly correlated (Supplementary Figure 1), their importance in the model might be underestimated. Nevertheless, we do not have evidence that these features negatively affect the prediction performance; hence, we kept them in the model.

eQTG-Finder uses known QTGs (*i*.*e*., causal genes for a phenotype QTL) as positive instances for model training because of the limited number of known eQTGs. We argue that QTGs are relevant for prioritizing eQTG since variation at the molecular level (*e*.*g*., in gene expression, metabolite, or protein level) can be propagated and cause variation at higher phenotypic levels (Fu et al. 2009; Civelek and Lusis 2013). For example, genetic variations in *AOP2* and *MAM1* cause *cis*-eQTLs for gene expression and metabolite QTLs for aliphatic glucosinolate biosynthesis, which confer insect resistance in Arabidopsis (Wentzell et al. 2007; Jansen et al. 2009). Both genes were prioritized in the top 5% rank percentile by eQTG-Finder. This result suggests that eQTG-Finder can identify QTLs for other molecular phenotypes, including metabolite and protein.

A lack of model interpretability may hamper a user’s comprehensive evaluation and assessment of the prioritization results. Regardless of the good performance, it is difficult to precisely understand how eQTG-Finder classifies certain genes as causal and others as non-causal, a typical issue for a complex model like Random Forest (Petch et al. 2022). Instead, in AraQTL, we provide additional sources of evidence to support the eQTG-Finder prioritization results (Hartanto *et al*., unpublished). For example, eQTG-Finder prioritizes transcription factor *LHY* as the regulator of *GLK2* (Figure 5). The network visualization in AraQTL showed that *LHY* is connected to *GLK2* by transcription factor binding site evidence, indicating that *LHY* may bind to the *GLK2* promoter and modulate its expression. Incorporating eQTG-Finder in the AraQTL web interface facilitates researchers to identify key regulators for genes of interest without the need for computational skills.

In the independent validation, some eQTG candidates were ranked poorly by eQTL-Finder (Figure 3). Low ranked assumed eQTG genes from the hypothetical and hypothetical-orthologs groups might not be actual eQTGs; however, the strong eQTG candidate ELF3 was also ranked poorly by both eQTG-Finder (61.2%) and QTG-Finder (44%). *ELF3* encodes a nuclear protein and was demonstrated to regulate gene expression leading to shade-avoidance response (Jimenez-Gomez et al. 2010). The complexity of the eQTG-Finder algorithm makes it difficult to dissect the prediction for *ELF3*. We investigated two of the most important features and noticed that this gene only has one identified protein domain and one paralog copy number, which is lower than the median values of causal genes (four and seventeen, respectively).

Likely, some features associated with eQTG are still missing in our model or underrepresented in our set of positive instances. Since the regulator-target relationship is specific, we expect that features representing gene-gene/protein-protein relationships (for example, STRING scores (Szklarczyk et al. 2019), transcription factor binding sites (Tian et al. 2020), and gene ontology semantic similarity (Yu 2020)) are relevant for prioritizing eQTG. Including these would shift the prioritization of generic eQTGs based on gene properties to the prioritization of eQTGs for a specific target using features based on gene-pair relationships. This is similar to the approaches of Wong et al. (2004) and Pandey et al. (2010), who predicted genetic interaction using gene pair relationships in yeast. The number of positive examples (*i*.*e*., confirmed eQTG-target pairs) is currently too small to properly train such a model for Arabidopsis. However, as data regarding genetic regulation is steadily increasing, we are optimistic that this strategy will be possible in the future.

## Data availability

The code and data for the analysis and visualization is available at the Wageningen University GitLab repository (https://git.wur.nl/harta003/eqtg-finder). eQTG-Finder prioritization is available at AraQTL (https://www.bioinformatics.nl/AraQTL/; Nijveen et al. 2017)

## Acknowledgments

We thank members of the Bioinformatics Group, Wageningen University, for feedback and suggestions.

## Conflict of Interest

We declare no conflict of interest.

## SUPPLEMENTARY FIGURES

**Supplementary Figure 1.**
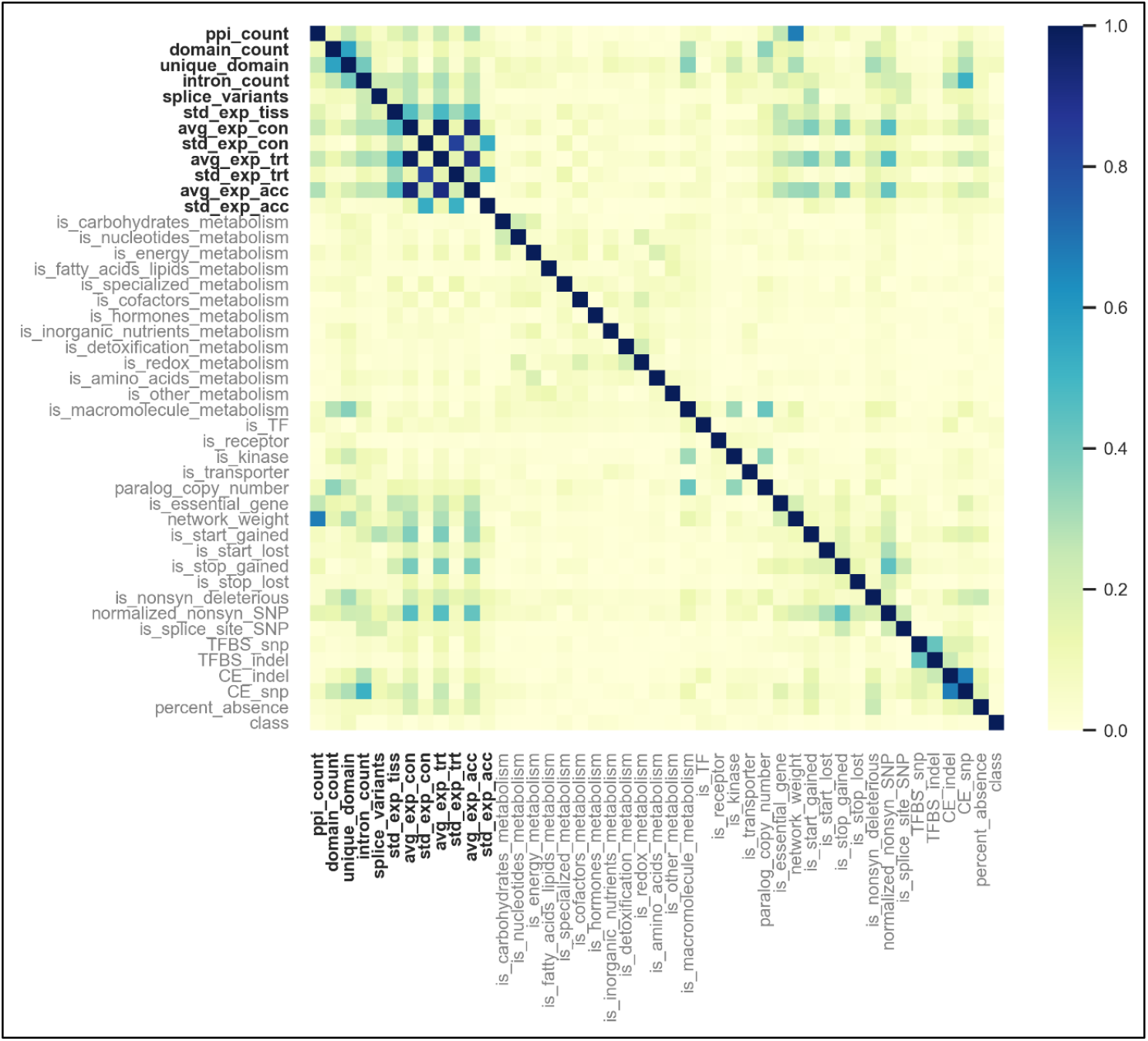
Correlation matrix of features used in the machine learning model. New features are indicated in bold.

**Supplementary Figure 2.**
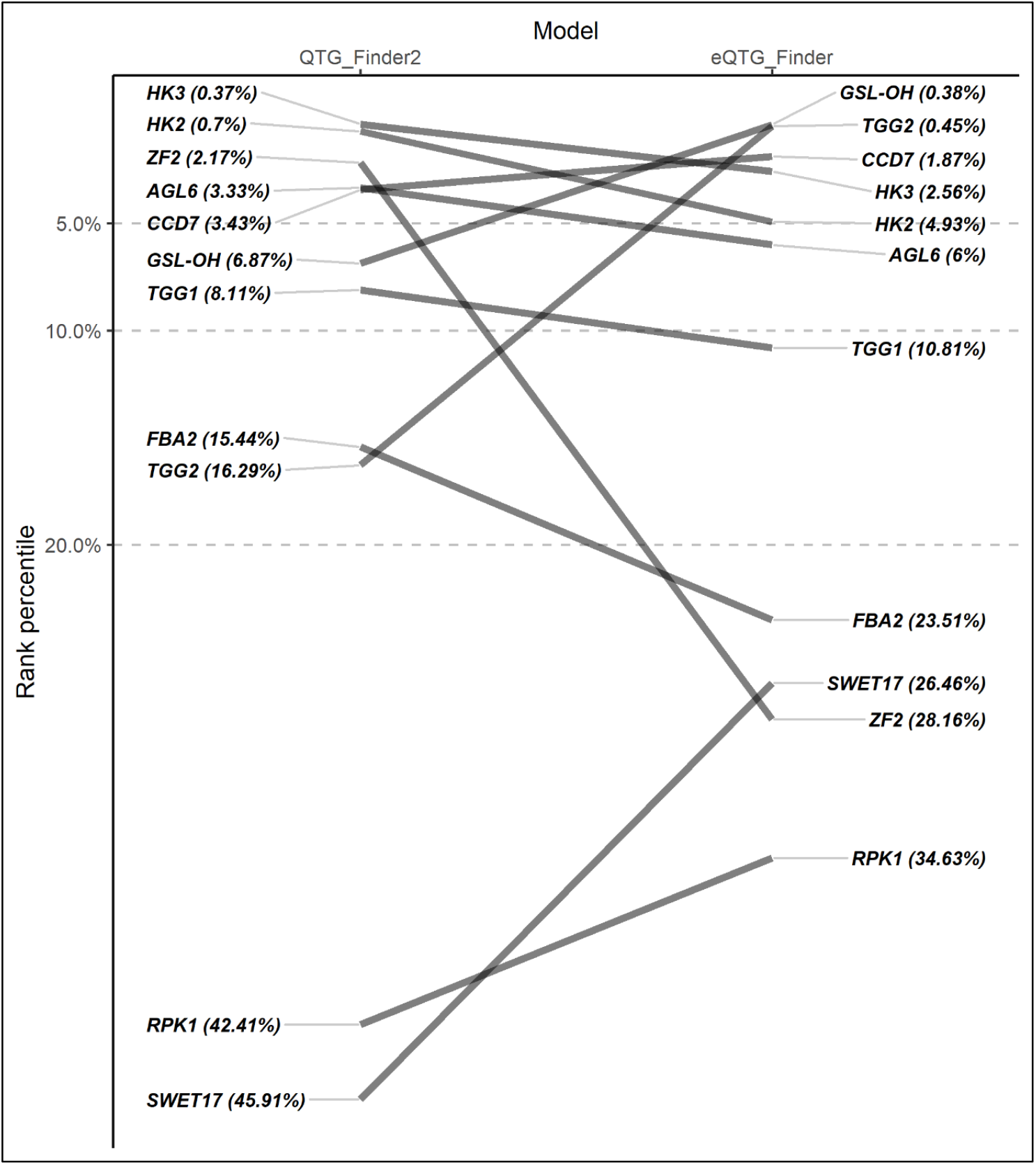
Rank percentile comparison of eleven original validation gene sets using the model with new features and the original model (QTG-Finder2).

## SUPPLEMENTARY TABLES

**Supplementary Table 2.**
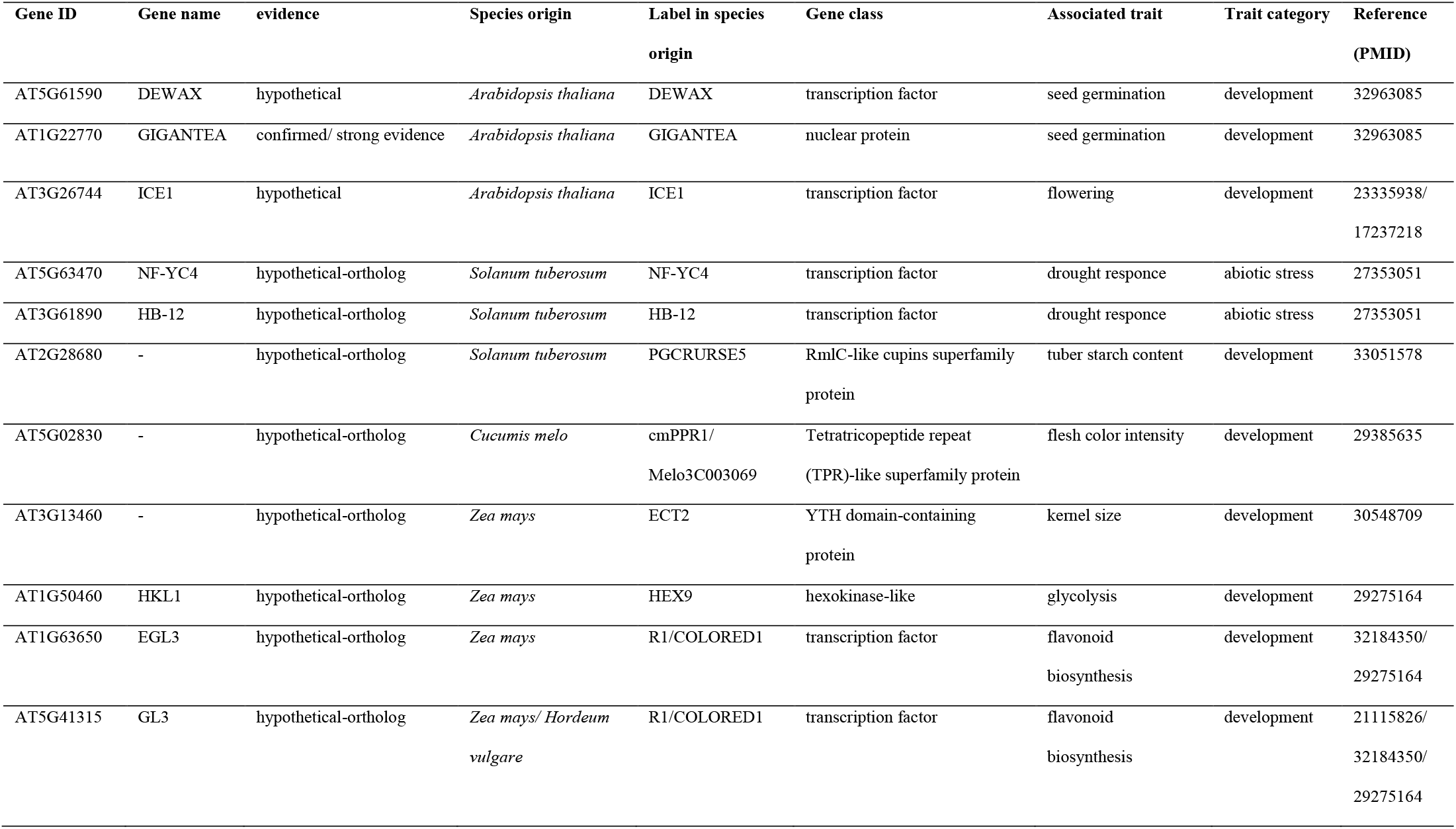
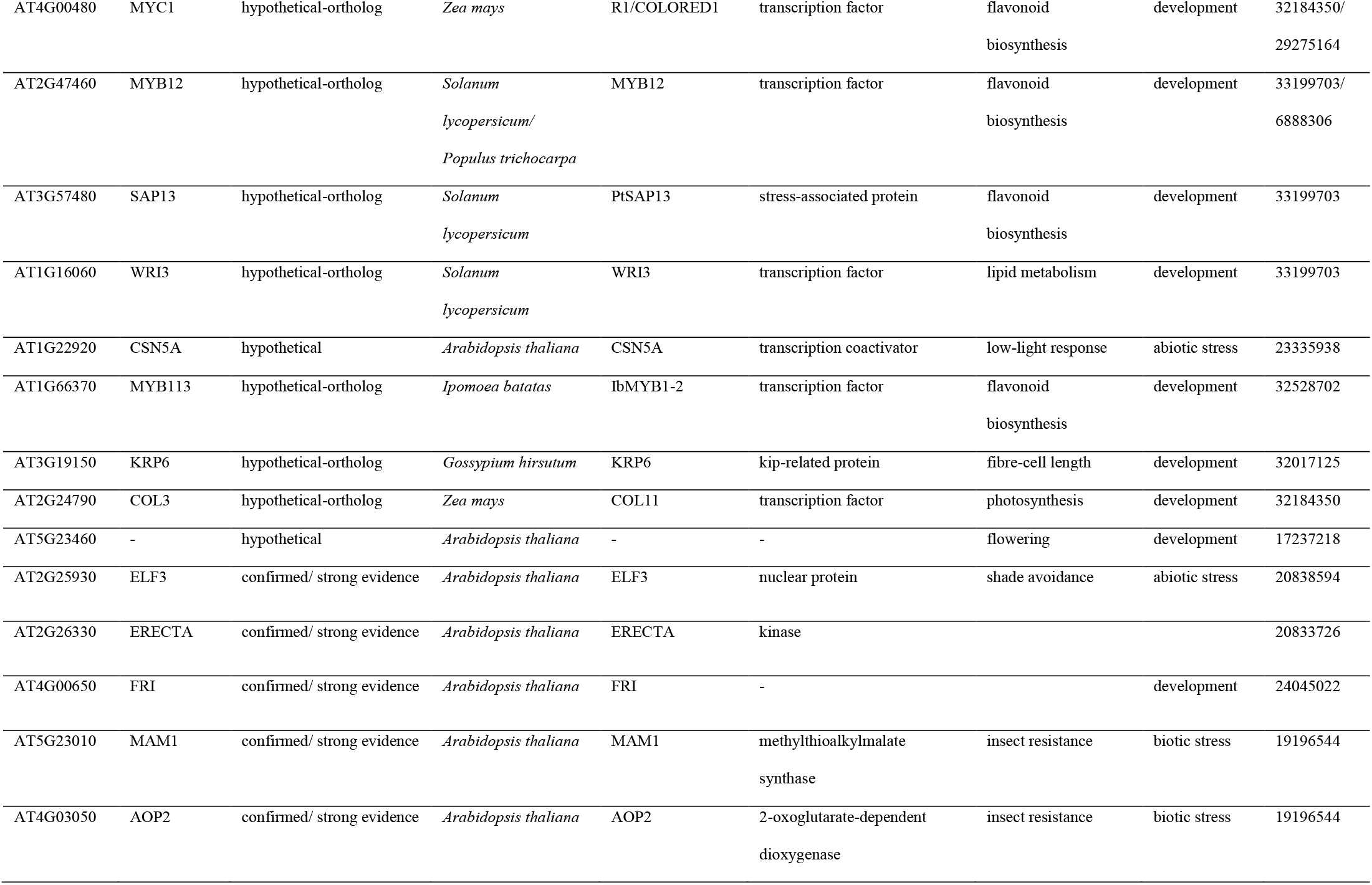
The list of candidate eQTGs used for independent validation

**Supplementary Table 3.** Wilcoxon Rank Sum Test statistics and p values for the difference in median of new features between causal genes and the other genes in the genome.

| **W** |  |  |
| --- | --- | --- |
| **Feature** | **statistics** | **p-value** |
| std exp tiss | 3312857 | $2.51 \times 10^8$ |
| avg exp con | 2624347 | 0.51 |
| std exp con | 3207362 | $3.86 \times 10^6$ |
| avg exp trt | 2531055 | 0.13 |
| std exp trt | 3354636 | $2.64 \times 10^9$ |
| avg exp acc | 2445143 | 0.02 |
| std exp acc | 3287936 | $8.19 \times 10^8$ |
| intron count | 2937908 | 0.03 |
| splice variants | 2805881 | 0.17 |
| ppi count | 3142925 | $3.75 \times 10^5$ |
| domain count | 3705449 | $3 \times 10^{20}$ |
| unique domain | 3944166 | $3.12 \times 10^{30}$ |

**Supplementary Table 4.**
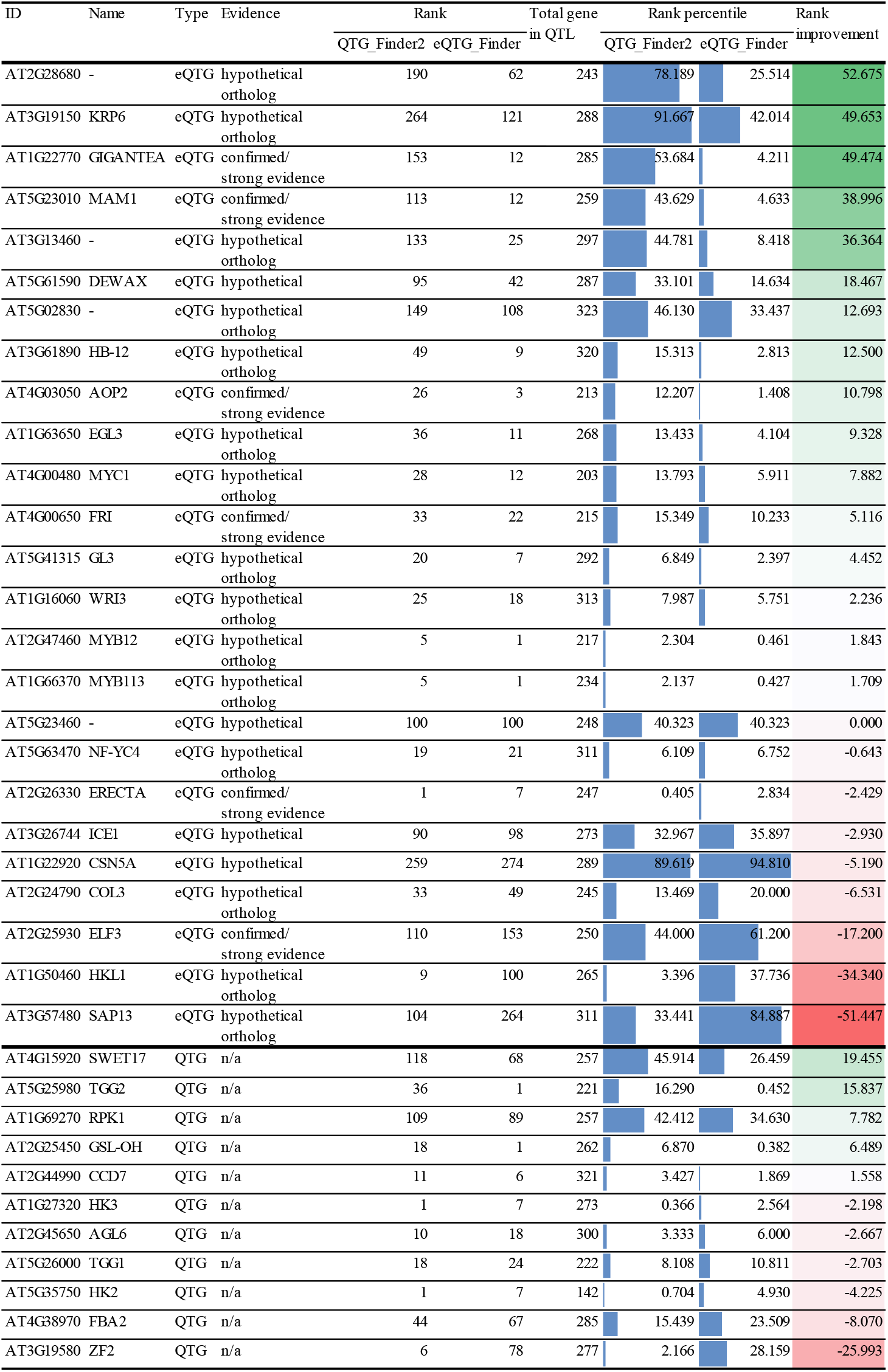
Candidate eQTL genes and their rank percentile based on the original QTG-Finder2 and eQTG-Finder.

**Supplementary Table 5.** Gene ontology terms enriched in the top 5% genes predicted as causal.

| GO ID | Term | Annotated | Significant | Expected | p-value | FDR |
| --- | --- | --- | --- | --- | --- | --- |
| GO:0030154 | cell differentiation | 675 | 118 | 37.74 | $7.50 \times 10^{30}$ | $4.51 \times 10^{26}$ |
| GO:0006468 | protein phosphorylation | 1037 | 141 | 57.98 | $3.00 \times 10^{22}$ | $9.02 \times 10^{19}$ |
| GO:0051762 | sesquiterpene biosynthetic<br>process | 25 | 19 | 1.4 | $1.80 \times 10^{19}$ | $3.61 \times 10^{16}$ |
| GO:0006355 | regulation of transcription,<br>DNA-templat... | 2053 | 279 | 114.78 | $2.60 \times 10^{19}$ | $3.91 \times 10^{16}$ |
| GO:0007165 | signal transduction | 1429 | 167 | 79.89 | $7.80 \times 10^{19}$ | $9.38 \times 10^{16}$ |
| GO:0048544 | recognition of pollen | 43 | 23 | 2.4 | $4.20 \times 10^{18}$ | $4.21 \times 10^{15}$ |
| GO:0045893 | positive regulation of<br>transcription, DN... | 509 | 96 | 28.46 | $1.40 \times 10^{16}$ | $1.20 \times 10^{13}$ |
| GO:0042742 | defense response to<br>bacterium | 430 | 69 | 24.04 | $1.70 \times 10^{14}$ | $1.28 \times 10^{11}$ |
| GO:0009686 | gibberellin biosynthetic<br>process | 29 | 14 | 1.62 | $3.00 \times 10^{11}$ | $2.01 \times 10^8$ |
| GO:0070588 | calcium ion transmembrane<br>transport | 45 | 16 | 2.52 | $2.70 \times 10^{10}$ | $1.62 \times 10^7$ |
| GO:0050832 | defense response to fungus | 257 | 41 | 14.37 | $3.20 \times 10^{10}$ | $1.63 \times 10^7$ |
| GO:0009753 | response to jasmonic acid | 189 | 37 | 10.57 | $3.70 \times 10^{10}$ | $1.63 \times 10^7$ |
| GO:0009611 | response to wounding | 215 | 37 | 12.02 | $3.70 \times 10^{10}$ | $1.63 \times 10^7$ |
| GO:0016114 | terpenoid biosynthetic<br>process | 146 | 49 | 8.16 | $3.80 \times 10^{10}$ | $1.63 \times 10^7$ |
| GO:0061408 | positive regulation of<br>transcription fro... | 24 | 12 | 1.34 | $1.30 \times 10^9$ | $5.21 \times 10^7$ |
| GO:0009414 | response to water<br>deprivation | 381 | 52 | 21.3 | $1.50 \times 10^9$ | $5.64 \times 10^7$ |
| GO:0045087 | innate immune response | 348 | 55 | 19.46 | $7.00 \times 10^9$ | $2.48 \times 10^6$ |
| GO:0009617 | response to bacterium | 508 | 87 | 28.4 | $2.90 \times 10^8$ | $9.69 \times 10^6$ |
| GO:0006952 | defense response | 1046 | 166 | 58.48 | $4.80 \times 10^8$ | $1.52 \times 10^5$ |
| GO:2000652 | regulation of secondary cell<br>wall biogen... | 28 | 13 | 1.57 | $5.60 \times 10^8$ | $1.68 \times 10^5$ |
| GO:0045490 | pectin catabolic process | 96 | 21 | 5.37 | $5.90 \times 10^8$ | $1.69 \times 10^5$ |
| GO:0002229 | defense response to<br>oomycetes | 76 | 18 | 4.25 | 6.60 x 10<sup>8</sup> | 1.78 x 10<sup>5</sup> |
| GO:0010200 | response to chitin | 141 | 26 | 7.88 | 6.80 x 10<sup>8</sup> | 1.78 x 10<sup>5</sup> |
| GO:0046777 | protein autophosphorylation | 191 | 31 | 10.68 | 8.40 x 10<sup>8</sup> | 2.11 x 10<sup>5</sup> |
| GO:0045487 | gibberellin catabolic process | 7 | 6 | 0.39 | 2.00 x 10<sup>7</sup> | 4.81 x 10<sup>5</sup> |
| GO:0042545 | cell wall modification | 168 | 33 | 9.39 | 2.40 x 10<sup>7</sup> | 5.35 x 10<sup>5</sup> |
| GO:0045944 | positive regulation of<br>transcription by ... | 220 | 42 | 12.3 | 2.40 x 10<sup>7</sup> | 5.35 x 10<sup>5</sup> |
| GO:0009735 | response to cytokinin | 104 | 20 | 5.81 | 2.60 x 10<sup>7</sup> | 5.59 x 10<sup>5</sup> |
| GO:0080027 | response to herbivore | 15 | 8 | 0.84 | 4.20 x 10<sup>7</sup> | 8.71 x 10<sup>5</sup> |
| GO:0019264 | glycine biosynthetic process<br>from serine | 5 | 5 | 0.28 | 5.40 x 10<sup>7</sup> | 0.0001 |
| GO:1904482 | cellular response to<br>tetrahydrofolate | 5 | 5 | 0.28 | 5.40 x 10<sup>7</sup> | 0.0001 |
| GO:0006565 | L-serine catabolic process | 5 | 5 | 0.28 | 5.40 x 10<sup>7</sup> | 0.0001 |
| GO:0045892 | negative regulation of<br>transcription, DN... | 296 | 35 | 16.55 | 1.00 x 10<sup>6</sup> | 0.0002 |
| GO:0048481 | plant ovule development | 52 | 15 | 2.91 | 2.00 x 10<sup>6</sup> | 0.0004 |
| GO:0009625 | response to insect | 30 | 10 | 1.68 | 3.10 x 10<sup>6</sup> | 0.0005 |
| GO:0046655 | folic acid metabolic process | 6 | 5 | 0.34 | 3.10 x 10<sup>6</sup> | 0.0005 |
| GO:0009416 | response to light stimulus | 741 | 90 | 41.43 | 7.80 x 10<sup>6</sup> | 0.001 |
| GO:0009809 | lignin biosynthetic process | 49 | 13 | 2.74 | 8.10 x 10<sup>6</sup> | 0.001 |
| GO:0007166 | cell surface receptor<br>signaling pathway | 49 | 13 | 2.74 | 9.30 x 10<sup>6</sup> | 0.001 |
| GO:0010114 | response to red light | 60 | 13 | 3.35 | 1.50 x 10<sup>5</sup> | 0.002 |
| GO:0009620 | response to fungus | 327 | 57 | 18.28 | 1.80 x 10<sup>5</sup> | 0.003 |
| GO:0005983 | starch catabolic process | 17 | 7 | 0.95 | 2.00 x 10<sup>5</sup> | 0.003 |
| GO:0010093 | specification of floral organ<br>identity | 13 | 7 | 0.73 | 2.10 x 10<sup>5</sup> | 0.003 |
| GO:0010951 | negative regulation of<br>endopeptidase act... | 12 | 6 | 0.67 | 2.10 x 10<sup>5</sup> | 0.003 |
| GO:0009944 | polarity specification of<br>adaxial/abaxia... | 23 | 8 | 1.29 | 2.10 x 10<sup>5</sup> | 0.003 |
| GO:0009828 | plant-type cell wall<br>loosening | 37 | 10 | 2.07 | 2.50 x 10<sup>5</sup> | 0.003 |
| GO:0009693 | ethylene biosynthetic<br>process | 30 | 9 | 1.68 | 2.50 x 10<sup>5</sup> | 0.003 |
| GO:0006979 | response to oxidative stress | 453 | 57 | 25.33 | 4.30 x 10<sup>5</sup> | 0.005 |
| GO:0055114 | oxidation-reduction process | 645 | 63 | 36.06 | 4.40 x 10<sup>5</sup> | 0.005 |
| GO:0030574 | collagen catabolic process | 5 | 4 | 0.28 | 4.60 x 10<sup>5</sup> | 0.006 |
| GO:0000266 | mitochondrial fission | 14 | 6 | 0.78 | 6.10 x 10<sup>5</sup> | 0.007 |
| GO:0010087 | phloem or xylem<br>histogenesis | 101 | 19 | 5.65 | 9.80 x 10<sup>5</sup> | 0.011 |
| GO:0080086 | stamen filament<br>development | 10 | 5 | 0.56 | 0.0001 | 0.012 |
| GO:0070370 | cellular heat acclimation | 10 | 5 | 0.56 | 0.0001 | 0.012 |
| GO:0009957 | epidermal cell fate<br>specification | 6 | 4 | 0.34 | 0.0001 | 0.014 |
| GO:0010106 | cellular response to iron ion<br>starvation | 6 | 4 | 0.34 | 0.0001 | 0.014 |
| GO:0009651 | response to salt stress | 461 | 47 | 25.77 | 0.0002 | 0.016 |
| GO:0097054 | L-glutamate biosynthetic<br>process | 3 | 3 | 0.17 | 0.0002 | 0.017 |
| GO:0016099 | monoterpenoid biosynthetic<br>process | 3 | 3 | 0.17 | 0.0002 | 0.017 |
| GO:0009823 | cytokinin catabolic process | 3 | 3 | 0.17 | 0.0002 | 0.017 |
| GO:1900386 | positive regulation of<br>flavonol biosynth... | 3 | 3 | 0.17 | 0.0002 | 0.017 |
| GO:0010311 | lateral root formation | 55 | 11 | 3.07 | 0.0002 | 0.019 |
| GO:0031408 | oxylipin biosynthetic<br>process | 17 | 6 | 0.95 | 0.0002 | 0.021 |
| GO:0051301 | cell division | 258 | 26 | 14.42 | 0.0003 | 0.025 |
| GO:0006826 | iron ion transport | 63 | 12 | 3.52 | 0.0003 | 0.03 |
| GO:0009737 | response to abscisic acid | 541 | 55 | 30.25 | 0.0004 | 0.032 |
| GO:0055072 | iron ion homeostasis | 93 | 16 | 5.2 | 0.0005 | 0.047 |

## Notes

https://git.wur.nl/harta003/eqtg-finder

### Competing Interest Statement

The authors have declared no competing interest.

