## Supplementary figures and images for "Prioritizing Candidate eQTL Causal Genes in Arabidopsis using Random Forests"

### Supplementary Figure 1

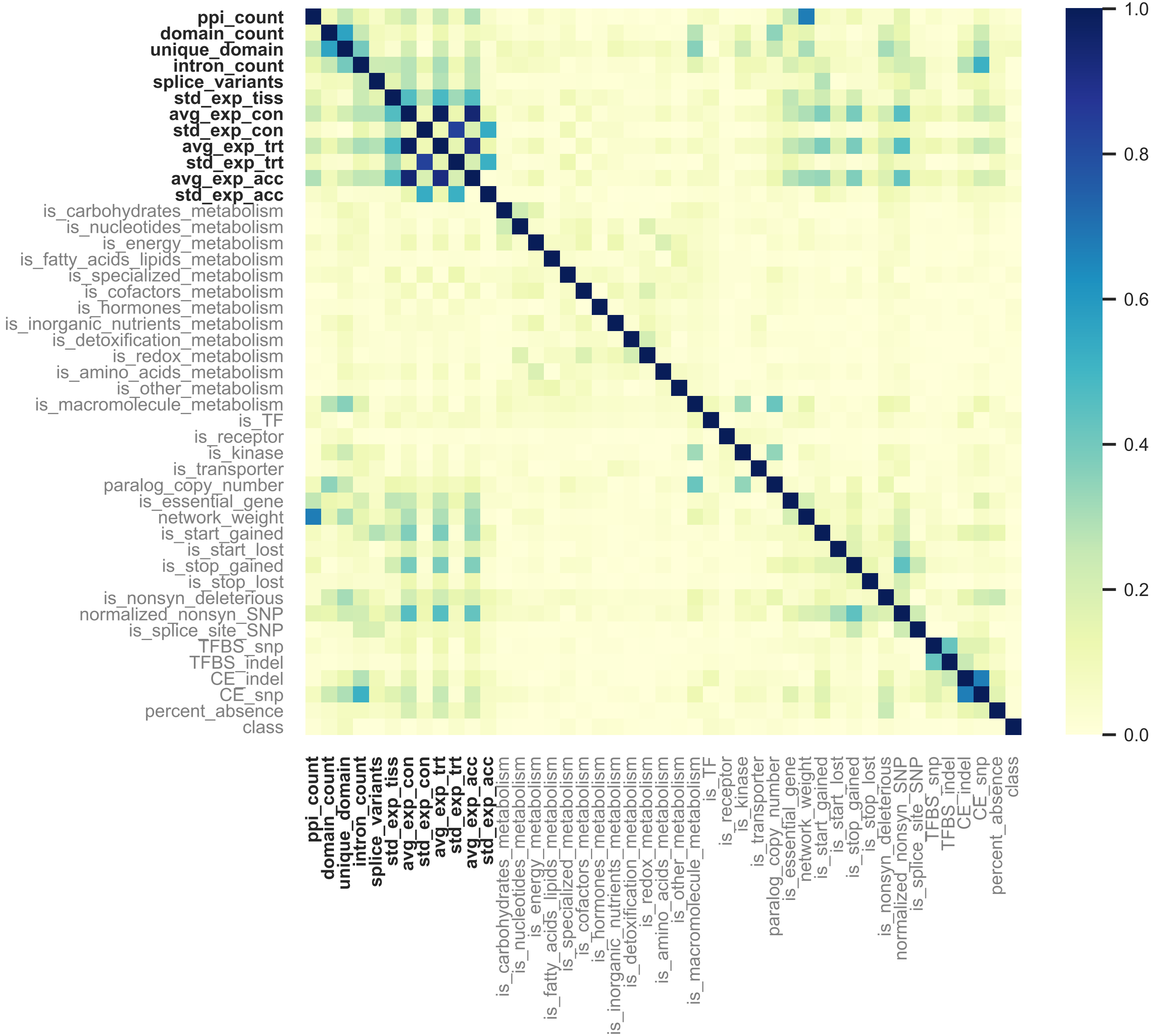

### Supplementary Figure 2

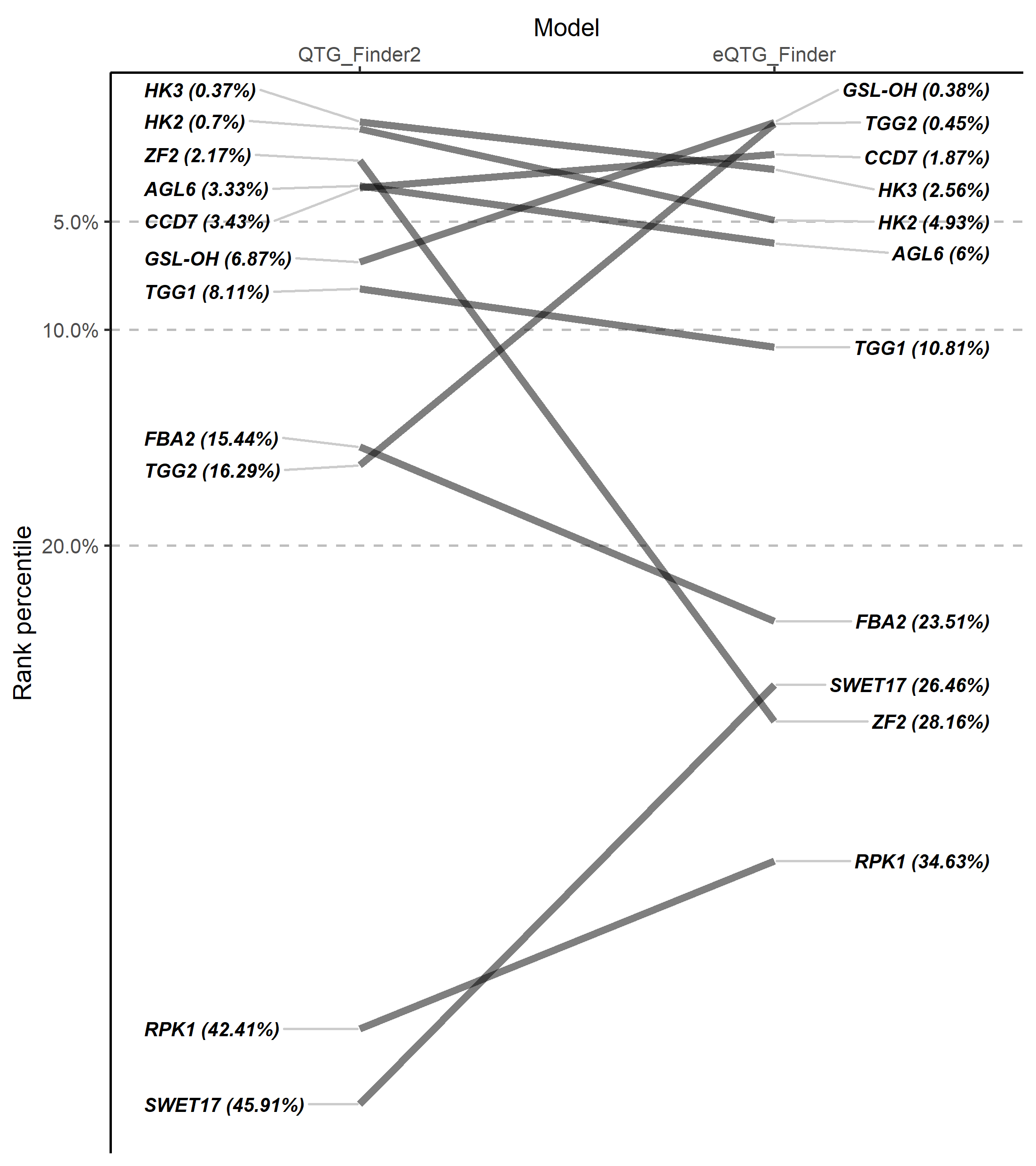
